# Elephant seal muscle cells adapt to sustained glucocorticoid exposure by shifting their metabolic phenotype

**DOI:** 10.1101/2021.02.05.429992

**Authors:** Julia Maria Torres-Velarde, Sree Rohit Raj Kolora, Jane I. Khudyakov, Daniel E. Crocker, Peter H. Sudmant, José Pablo Vázquez-Medina

**Author notes:** Correspondence: José Pablo Vázquez-Medina Department of Integrative Biology University of California, Berkeley, 3040 Valley Life Sciences Bldg. #3140, Berkeley CA 94720-3140.

## Abstract

Elephant seals experience natural periods of prolonged food deprivation while breeding, molting, and undergoing postnatal development. Prolonged food deprivation in elephant seals increases circulating glucocorticoids without inducing muscle atrophy, but the cellular mechanisms that allow elephant seals to cope with such conditions remain elusive. We generated a cellular model and conducted transcriptomic, metabolic, and morphological analyses to study how seal cells adapt to sustained glucocorticoid exposure. Seal muscle progenitor cells differentiate into contractile myotubes with a distinctive morphology, gene expression profile, and metabolic phenotype. Exposure to dexamethasone at three ascending concentrations for 48h modulated the expression of 6 clusters of genes related to structural constituents of muscle and pathways associated with energy metabolism and cell survival. Knockdown of the glucocorticoid receptor (GR) and downstream expression analyses corroborated that GR mediates the observed effects. Dexamethasone also decreased cellular respiration, shifted the metabolic phenotype towards glycolysis, and induced mitochondrial fission and dissociation of mitochondria-ER interactions without decreasing cell viability. Knockdown of DDIT4, a GR target involved in the dissociation of mitochondria-ER membranes, recovered respiration and modulated antioxidant gene expression. These results show that adaptation to sustained glucocorticoid exposure in elephant seal myotubes involves a metabolic shift toward glycolysis, which is supported by alterations in mitochondrial morphology and a reduction in mitochondria-ER interactions, resulting in decreased respiration without compromising cell survival.

## Introduction

Glucocorticoids are steroid hormones produced and released by the adrenal cortex in response to circadian changes in metabolism and stress perturbations (Sapolsky, Romero, & Munck, 2000; Tsigos & Chrousos, 2002). Activation of the hypothalamic-pituitary-adrenal axis controls glucocorticoid secretion by releasing corticotropin releasing hormone (CRH) from the hypothalamus (Vale, Spiess, Rivier, & Rivier, 1981). CRH then stimulates the synthesis and secretion of adrenocorticotropin (ACTH) from the anterior pituitary. ACTH acts on the adrenal cortex resulting in glucocorticoid and mineralocorticoid secretion (Chrousos, 1992).

Glucocorticoids participate in numerous physiological processes including metabolism, immune function, reproduction, growth, cognition and cardiovascular regulation (Baxter & Forsham, 1972; Cain & Cidlowski, 2017; Dickerson & Kemeny, 2004). The glucocorticoid receptor (GR), which belongs to the nuclear receptor family of ligand-dependent transcription factors (Evans, 1988; Luisi et al., 1991), mediates the biological actions of glucocorticoids by activating or repressing gene transcription (Oakley & Cidlowski, 2013). The cellular response to glucocorticoids is cell type-, tissue-, and species-specific (Lu & Cidlowski, 2006; Romero & Gormally, 2019). Similarly, acute and sustained exposure to glucocorticoids have contrasting biological effects. While acute exposure to glucocorticoids is adaptive, chronic exposure is considered deleterious (Sapolsky et al., 2000). In skeletal muscle, sustained exposure to glucocorticoids suppresses protein synthesis while inducing proteolysis and muscle mass loss, ultimately leading to muscle atrophy (Braun & Marks, 2015; Hasselgren, 1999; Schakman, Kalista, Barbé, Loumaye, & Thissen, 2013).

Northern elephant seals (*Mirounga angustirostris*) endure spontaneous long-term periods (2-3 months) of food deprivation while breeding, molting and undergoing postnatal development (Costa, Le Boeuf, Huntley, & Ortiz, 1986; C. L. Ortiz, Costa, & Le Boeuf, 1978; Reiter, Stinson, & Le Boeuf, 1978). During these natural prolonged fasting periods, elephant seals rely primarily on lipid metabolism and endogenous glucose production to support metabolism in glucose-dependent tissues (Champagne, Crocker, Fowler, & Houser, 2012; Crocker, Champagne, Fowler, & Houser, 2014). Notably, prolonged fasting in elephant seals increases the levels of circulating cortisol, the primary glucocorticoid in pinnipeds (Champagne, Houser, & Crocker, 2006; Jelincic, Tift, Houser, & Crocker, 2017; Khudyakov et al., 2019; R. M. Ortiz, Noren, Ortiz, & Talamantes, 2003; R. M. Ortiz, Wade, & Ortiz, 2001), without inducing muscle atrophy (Wright et al., 2020). The cellular mechanisms that drive tolerance to sustained glucocorticoid elevations in elephant seals, however, remain unexplored.

Recent work suggests that fasting-induced shifts in muscle metabolism stimulate pathways associated with preserving muscle mass in elephant seals (Wright et al., 2020), while earlier studies show that administration of exogenous ACTH upregulates catabolic genes in elephant seal skeletal muscle (Khudyakov, Champagne, Preeyanon, Ortiz, & Crocker, 2015; Khudyakov, Preeyanon, Champagne, Ortiz, & Crocker, 2015). Here, we generated a cellular model to study the mechanisms that drive tolerance to sustained glucocorticoid exposure in elephant seal muscle. We found that sustained glucocorticoid exposure promotes concerted changes in the expression of genes related to muscle remodeling and pathways associated with energy metabolism and cell survival in elephant seal myotubes. Furthermore, we found that adaptation to sustained glucocorticoid exposure in seal myotubes involves changes in mitochondrial morphology and function, and a shift in metabolic phenotype that promotes cell survival.

## Results

### Characterization of elephant seal muscle cells in primary culture

We developed a model to study the effects of sustained glucocorticoid exposure on muscle metabolism in elephant seals. Elephant seal myoblasts proliferate in primary culture and can be differentiated into spontaneously contracting myotubes (Figure 1A). Both myoblasts and myotubes stain positive for the muscle-specific protein desmin (Figure 1B) and express myogenic markers (Figure 1C). Oxygen consumption (Figure 1D) and cytochrome oxidase protein abundance (Figure 1E) are higher in myotubes than in myoblasts, suggesting increased mitochondrial content in myotubes compared to myoblasts. The total ATP production rate is twice as high in myotubes as in myoblasts (Figure 1F). The primary source of ATP in myoblasts is glycolysis while myotubes use glycolysis and respiration equally (Figure 1F). Therefore, while proliferating in primary culture, elephant seal myoblasts are predominantly glycolytic cells (glycolytic index = 72%, Figure 1F). In contrast, elephant seal myotubes rely on both glycolysis and respiration to produce ATP (glycolytic index = 54%, Figure 1F).

**Figure 1.**
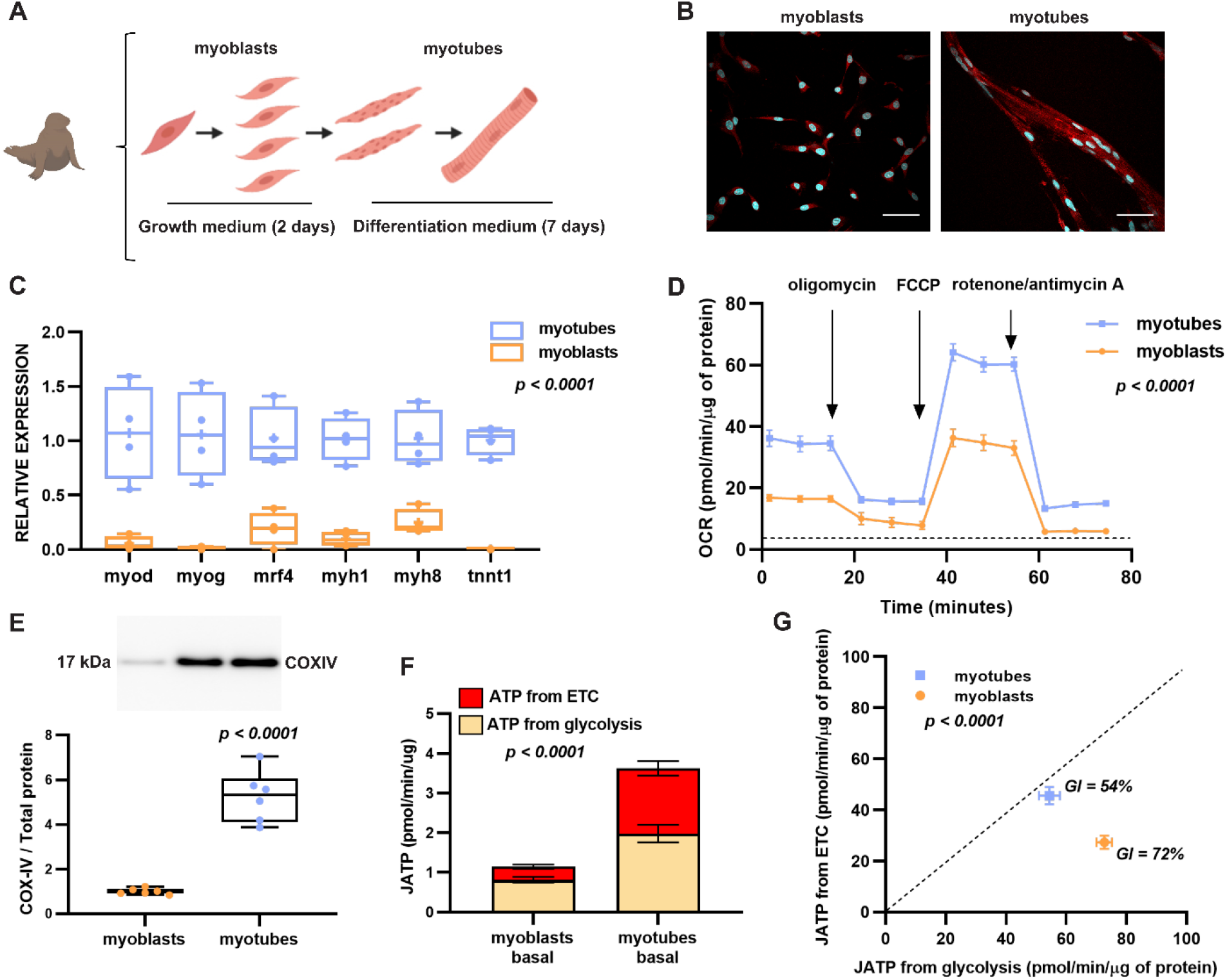
Characterization of elephant seal muscle cells in primary culture. A) Schematic representation of the protocol used to obtain elephant seal myotubes. B) Myoblasts and myotubes immunostained with desmin (red), scale bar is 33 μm. C) Myogenic gene expression in myoblasts vs myotubes, D) Oxygen consumption by myoblasts vs myotubes, E) Cytochrome oxidase protein abundance, F) Bioenergetic profile and G) glycolytic index (GI) of myoblasts and myotubes. Values were calculated according to (Mookerjee, Gerencser, Nicholls, & Brand, 2018). Results in D, F, and G are mean ± SEM, n=9.

### Sustained glucocorticoid exposure is associated with transcriptional signatures of muscle remodeling and cell survival

We incubated elephant seal myotubes with four ascending concentrations of the glucocorticoid dexamethasone (DEX) for 48 h and measured glucocorticoid (*gr*) and mineralocorticoid receptor (*mr*) expression using RT-qPCR. DEX decreased *gr* expression without affecting *mr* (Figure S1A). DEX also decreased GR protein abundance (Figure S1B). Co-treatment with the GR antagonist RU486 prevented the observed DEX-induced decrease in *gr* expression (Figure S1C). These results show that DEX signals mainly *via* GR in elephant seal myotubes.

We used RNA-seq to evaluate global changes in gene expression in myotubes incubated with DEX (Figure 2A) and identified six clusters of genes that changed in a concerted manner with increasing DEX doses (Figure 2B). We then conducted a Gene Ontology (GO) enrichment analysis of these clusters (Figure 2C). Cluster 3, which grouped genes that decreased with increasing DEX concentrations, was enriched for actin binding, cadherin binding, laminin binding, extracellular matrix structural constituents and insulin-like growth factor 1 binding. Cluster 5, which grouped genes that increased in the low and intermediate concentrations while decreasing in the highest dose, was enriched for structural constituents of muscle (Figure 2C). Cluster 6, which grouped genes upregulated with increasing DEX levels, was enriched for aspartic-type peptidase and endopeptidase activity. These results suggest that sustained exposure to glucocorticoids promotes muscle remodeling in elephant seal myotubes.

**Figure 2.**
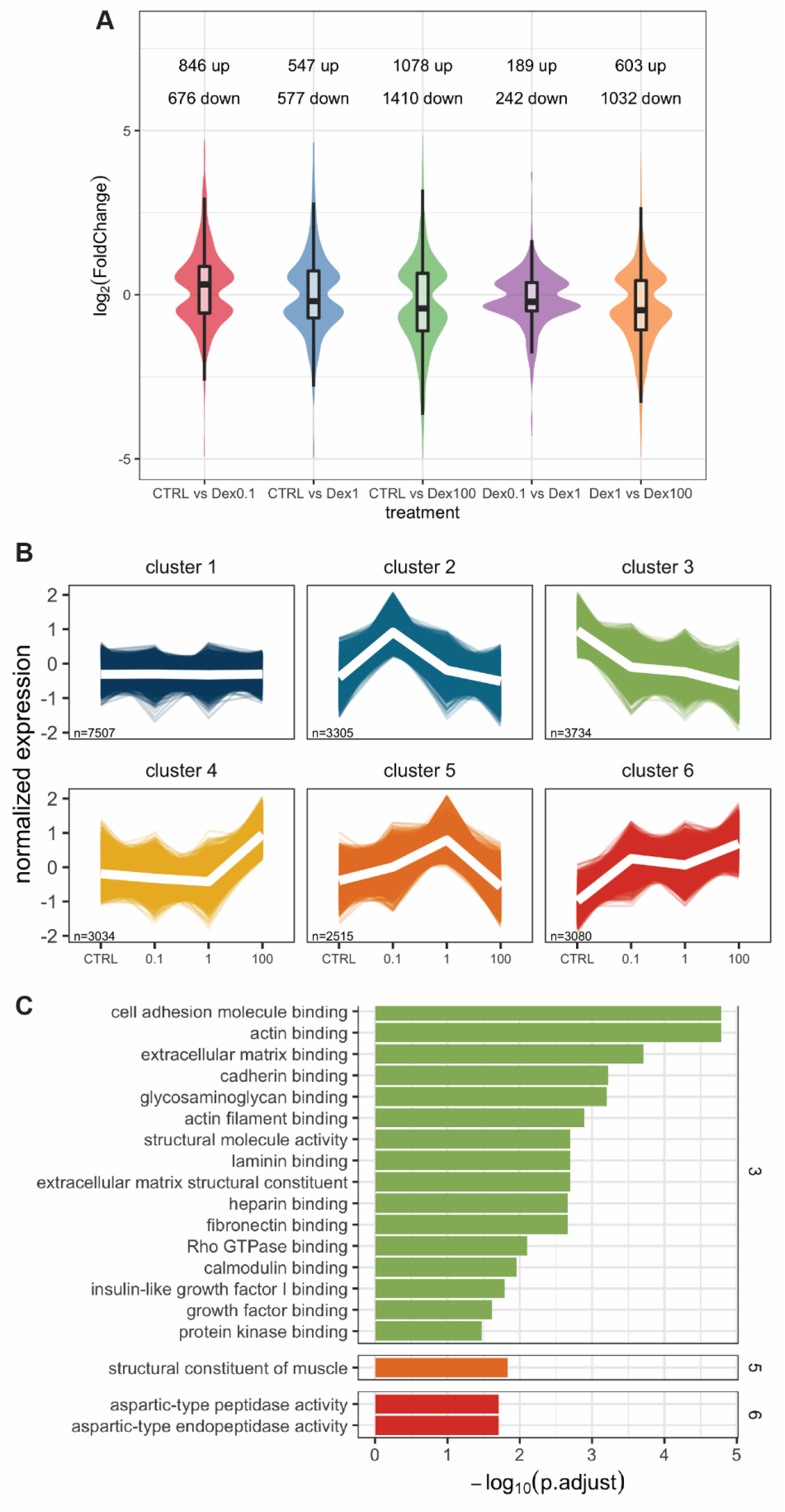
Transcriptome response to sustained glucocorticoid exposure in elephant seal myotubes. A) Differentially expressed genes in control vs DEX treatments and among DEX treatments. B) Gene expression clustering. C) GO terms for biological processes overrepresented in gene clusters.

We next identified genes differentially expressed (DE) in all DEX treatments (Figure 3A), and constructed functional interaction networks (FIN) based on enriched Reactome pathways (Wu, Feng, & Stein, 2010) (Figure 3B). We also generated FIN from genes DE in each condition compared to control (Figure S2). Pathways enriched at the low and intermediate DEX concentrations include the mechanistic target of rapamycin (mTOR) signaling, myogenesis, and protein digestion and absorption (Figure S2A, S2B), whereas pathways enriched only at the highest DEX concentration include pyruvate metabolism, autophagy, integration of energy metabolism, regulation of cholesterol biosynthesis by sterol regulatory element-binding protein (SREBP), and decreased striated muscle contraction (Figure S2C). In contrast, pathways enriched in all treatments include extracellular matrix organization, transforming growth factor-beta (TGF-β) signaling, hypoxia-inducible factor-1 alpha (HIF-1α) signaling, GR regulatory network, ras-related protein1 (Rap1) signaling, cadherin signaling and phosphoinositide 3-kinase (PI3K-Akt) signaling (Figure 3B). GO-cellular component analysis (Roncaglia et al., 2013) showed that focal adhesions, collagen-containing extracellular matrix, stress fibers, sarcolemma, cytoskeleton, and ER lumen are the major structures in which DE genes perform a biological function (data not shown).

**Figure 3.**
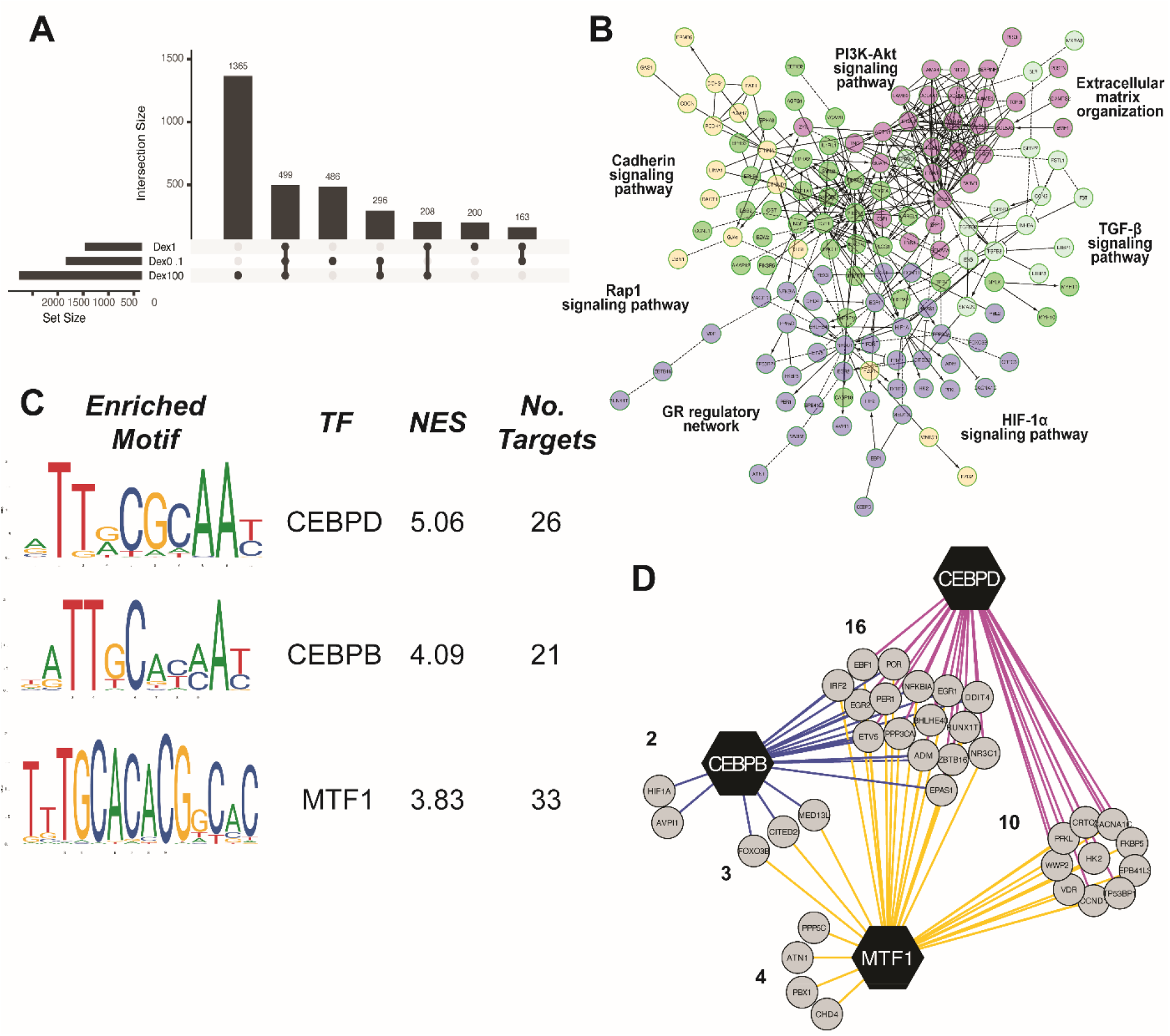
Functional Interaction Networks (FIN) and motif enrichment analysis in genes differentially expressed upon glucocorticoid treatment. A) UpSet plot of DE genes in myotubes treated with DEX. B) FIN built using Reactome pathways enriched in control vs all DEX treatments. Nodes represent genes, edges represent functional interactions, and different colors represent network modules. Lilac nodes represent genes in module 0. C) Motif enrichment analysis in the promoters of DE genes identified in module 0. TF: transcription factor, NES: normalized enrichment score. DNA-binding sites were obtained using JASPAR (Fornes et al., 2020). D) Regulatory network for transcription factors identified in module 0. CEBP: CCAAT/enhancer-binding protein, MTF1: metal regulatory transcription factor 1.

We then identified network modules that group genes DE in all treatments based on their biological process (Wu et al., 2010). We found 5 modules that grouped more than 15 genes (Figure 3B), including module 0, which grouped 40 genes. GO-cellular component analysis showed that proteins in module 0 are located primarily in the cytoplasm and the nucleus.

Pathways represented in module 0 include HIF-1□ signaling, circadian clock, and GR regulatory network (FDR < 0.05). We identified transcriptional regulators of genes in module 0 using cis-regulatory analysis (Janky et al., 2014) (Figure 3C) and constructed networks to identify genes regulated by these transcription factors (TFs) (Figure 3D). CEBPD, CEBPB and MTF1 were the major TFs identified in our analyses with 26, 21, and 33 direct targets respectively, and 16 shared targets (Figure 3D). CEBPD regulates the inflammatory response (Ko, Chang, & Wang, 2015), cell proliferation, differentiation, death and survival (Balamurugan & Sterneck, 2013). CEBPB maintains the undifferentiated state of satellite cells (Marchildon et al., 2012). MTF1, which is upregulated during myogenesis (Tavera-Montañez et al., 2019), modulates cellular adaptation to hypoxia (Murphy, Sato, Dalton, & Laderoute, 2005) and oxidative stress (Bahadorani, Mukai, Egli, & Hilliker, 2010). Similar to our results, previous work showed that *ddit4*, a known CEBPD target (Lin, Qian, Shi, & Chen, 2005) and one of the 16 shared genes identified in our cis-regulatory analysis, is one of the top upregulated genes in skeletal muscle of elephant seals undergoing acute ACTH stimulation (Khudyakov, Champagne, et al., 2015). Overall, our results show that sustained exposure to glucocorticoids is associated with transcriptional signatures of muscle remodeling and cell survival in elephant seal myotubes.

### Glucocorticoids regulate muscle remodeling via gr signaling

We confirmed the effect of sustained glucocorticoid exposure on the expression of specific genes involved in muscle remodeling using RT-qPCR. Consistent with the results of our cluster analyses, DEX had differential effects on myogenic genes. Expression of *myog* (*F=5*.*793, p=0*.*0014*) and *mrf4* (*F=71*.*83, p<0*.*0001*) increased with exposure to 0.1-10 µM DEX, while expression of all measured myogenic genes (Figure 4A), along with myosin heavy chain 1 protein levels (Figure 4B), decreased with exposure to the highest DEX concentration. Moreover, DEX increased *foxo1 (F= 6*.*031, p=0*.*0012), foxo3 (F=9*.*125, p=0*.*0009), cebpd (F= 66*.*91, p<0*.*0001)*, and *ddit4 (F=27*.*53, p<0*.*0001) expression*, suggesting activation of catabolic pathways (Figure 4A). In contrast, DEX did not increase the expression of genes associated with DEX-induced muscle wasting, including *murf1* and *myostatin* (Ma et al., 2003) (Figure S3). DEX-induced expression of catabolic genes was prevented by *gr* knockdown (Figure 4C, 4D). These results show that sustained exposure to glucocorticoids regulates muscle remodeling by suppressing myogenesis and upregulating catabolic genes *via* GR signaling in elephant seal myotubes.

**Figure 4.**
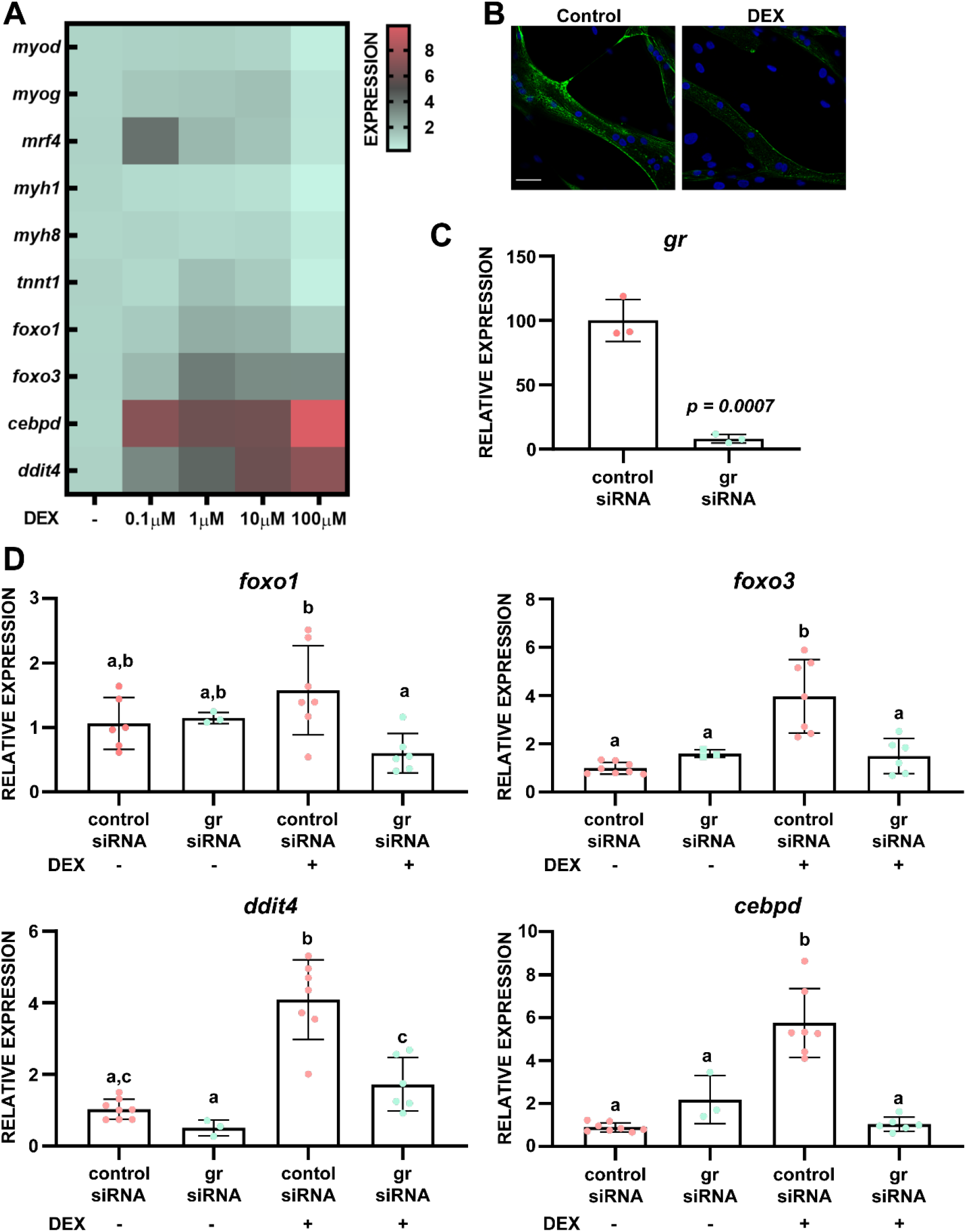
Glucocorticoids regulate muscle gene expression via GR signaling. A) mRNA levels of genes involved in muscle function and catabolism in myotubes treated with DEX for 48h, n=6. B) Myotubes treated with or without 100 µM DEX and immunostained for myosin heavy chain 1 (green), scale bar is 33 micrometers. C) gr knockdown using siRNA. D) Effect of gr knockdown on expression of foxo1 (F=4.468, p=0.0164), foxo3 (F=14.66, p<0.0001), ddit4 (F=27.66, p<0.0001) and cebpd (F=37.98, p<0.0001) in myotubes treated with or without 100 µM DEX. Different letters denote significant differences between treatments.

### Sustained glucocorticoid exposure promotes mitochondrial fission and suppresses mitochondrial respiration

We evaluated mitochondrial function in myotubes treated with DEX using extracellular flux assays. Basal respiration, maximal respiration, spare respiratory capacity, ATP-linked respiration, ETC-driven and total ATP production decreased with exposure to the higher DEX concentration (Figure 5A, 5B, 5C), suggesting overall suppression of mitochondrial metabolism. In contrast, DEX did not have an effect on glycolysis-driven ATP production (Figure 5C) or cell viability (Figure S4). Moreover, treatment with the lower DEX concentration had no effect on basal oxygen consumption, ATP-linked respiration, or total ATP production, but decreased maximal respiration and spare respiratory capacity (Figure 5A, 5B, 5C). Despite the observed changes in mitochondrial function, cells treated with the lower DEX concentration retained their respiratory and glycolytic capacity, whereas cells treated with the higher DEX dose became glycolytic (Figure 5E). Consistent with this observation, our RNA-seq analyses showed enrichment for the central carbon metabolism pathway and PI3K-Akt signaling, which are involved in the Warburg effect (Courtnay et al., 2015) (Figure S2C, Supplementary Table 3). Taken together, these results show that elephant seal myotubes respond to sustained glucocorticoid exposure by suppressing mitochondrial respiration and shifting their metabolic phenotype towards glycolysis.

**Figure 5.**
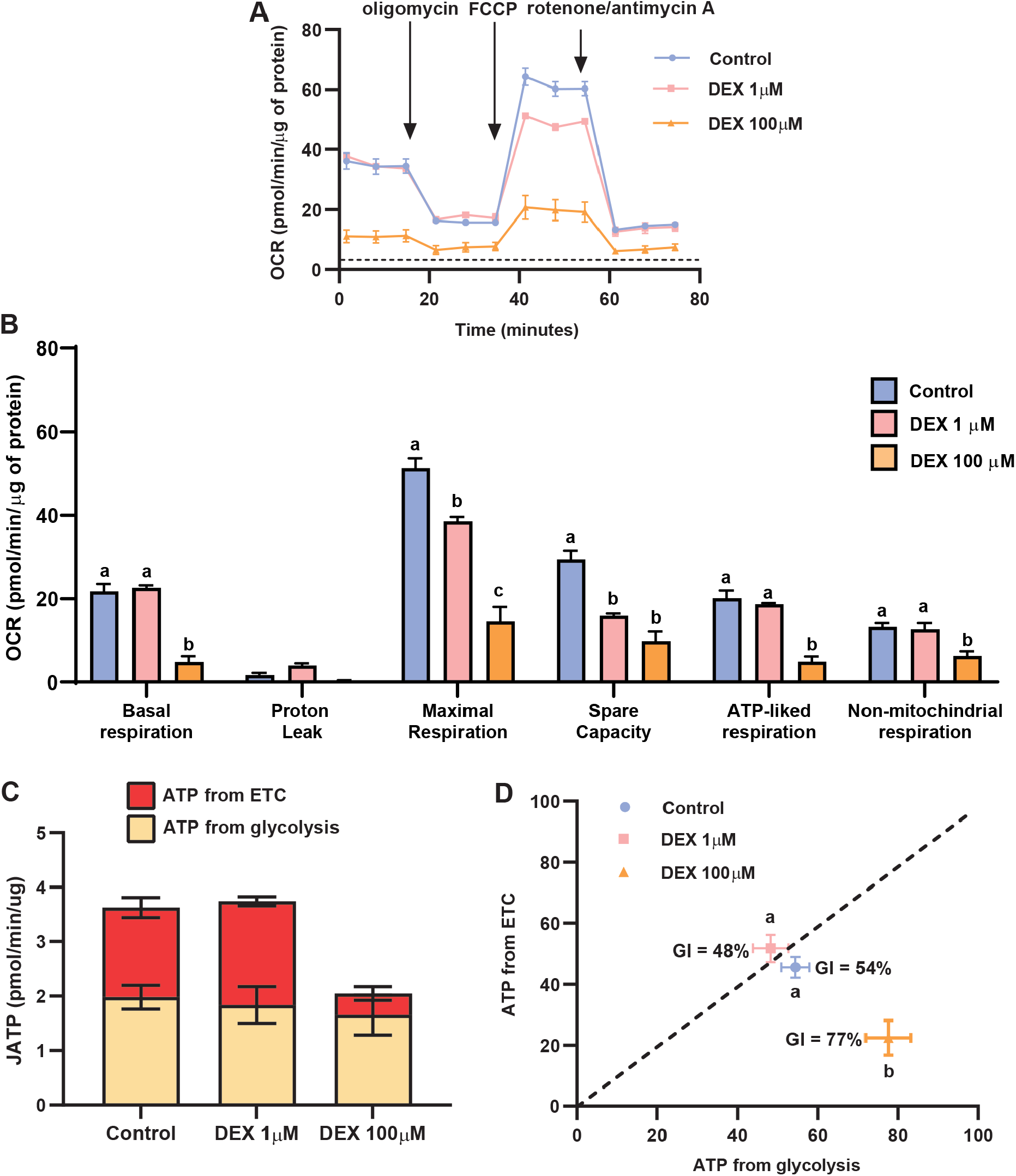
Sustained glucocorticoid exposure suppresses mitochondrial function. A) Effects of DEX exposure for 48h on oxygen consumption (F=6.747, p = 0.0035), B) mitochondrial function (F=50.42, p<0.0001), C) bioenergetic profile (F=5.453, p=0.0103) and D) glycolytic index (F=8.357, p=0.0015) of myotubes treated with or without 1 or 100 µM DEX. Results are expressed as mean ± SEM, n = 6. Different letters denote significant differences between treatments.

We next tested whether the observed effects of sustained glucocorticoid exposure on mitochondrial function are related to changes in mitochondrial morphology using 3D confocal microscopy (Figure 6A). DEX decreased the average length and degree of the mitochondrial network while increasing the total number of mitochondrial nodes, edges, connected components and free-end nodes suggesting a decrease in mitochondrial connections (Figure 6B) (Harwig et al., 2018). Similarly, the connectivity score decreased in cells treated with DEX, suggesting increased mitochondrial fragmentation (Figure 6C). DEX also decreased mitochondrial volume by 45%, total mitochondrial length by 45-48% (Figure 6C), and cytochrome oxidase abundance by 40% (Figure 6D). Moreover, DEX increased dynamin-related protein 1 (DRP1) phosphorylation (Figure 6E), which is essential for mitochondrial fission (Fonseca, Sánchez-Guerrero, Milosevic, & Raimundo, 2019). Overall, these results show that sustained exposure to glucocorticoids promotes mitochondrial fission in elephant seal myotubes. Furthermore, these results suggest that the observed changes in mitochondrial function are related to changes in the mitochondrial reticulum.

**Figure 6.**
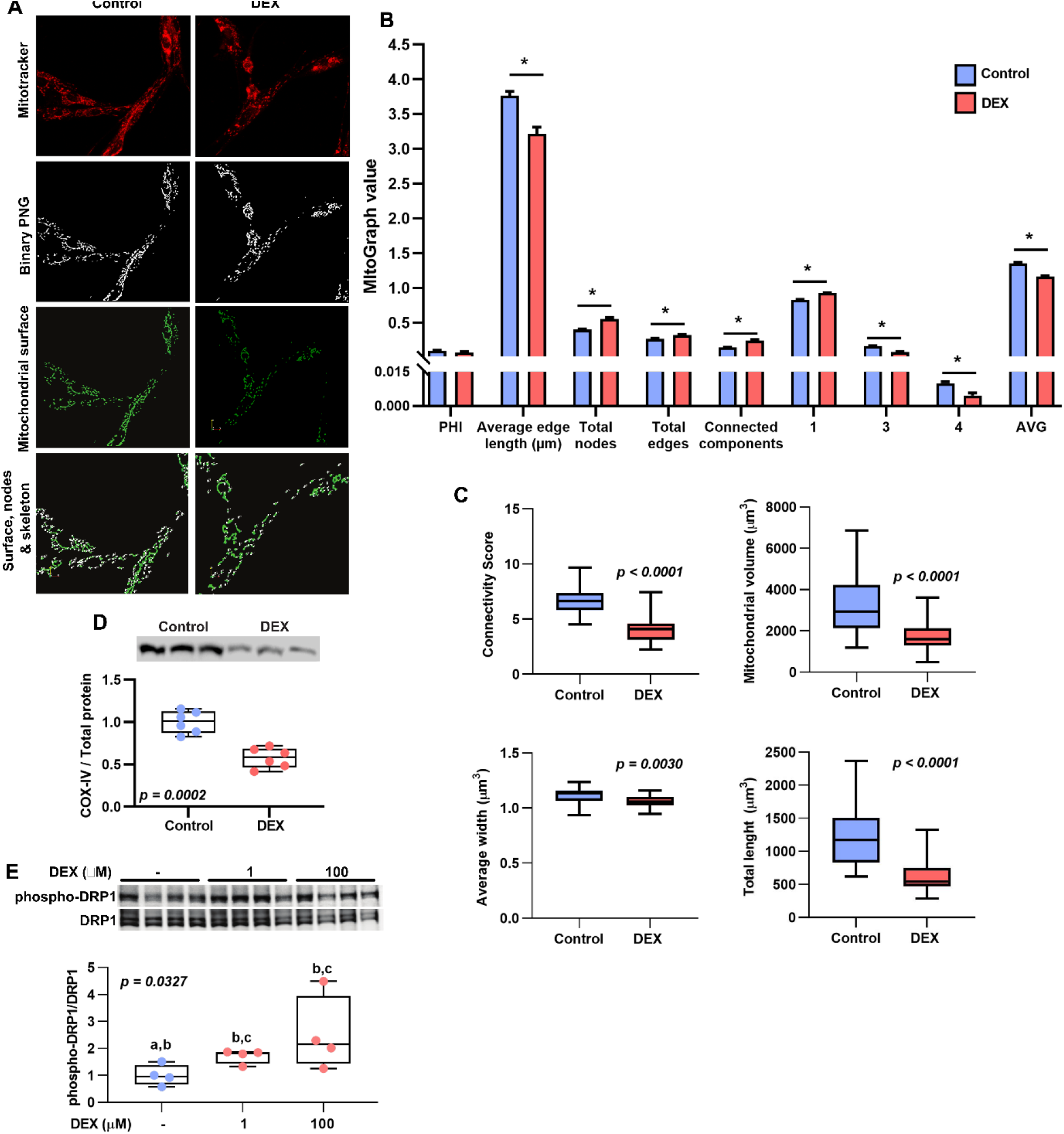
Sustained glucocorticoid exposure promotes mitochondrial fission. A) Myotubes treated with or without 100 µM DEX for 48h. Cells labeled with MitoTracker Red and Mitograph output files. B) Effects of DEX on the mitochondrial network. PHI: ratio of the largest connected component. Total nodes, edges, and number of connected components are normalized to total mitochondrial length to account for differences in myotube size. Numbers (degree distribution) indicate: 1 = free ends, 3 = three junctions, 4 = four junctions. AVG (average degree) represents the sum of each ratio (1, 3 or 4) multiplied by the corresponding number factor (1, 3 or 4). * = p < 0.001. C) Effects of DEX on mitochondrial connectivity score, volume, width and total length. Results represent at least 25 myotubes imaged in three replicates per treatment. D) COX-IV protein abundance and E) Dynamin-related protein 1 (DRP1) phosphorylation in myotubes treated with or without DEX (F=5.172, p=0.0320). Different letters denote significant differences between treatments. Whiskers are 1-99 percentile.

### Sustained glucocorticoid exposure promotes the dissociation of mitochondria-associated ER membranes

Mitochondria-ER interactions support cellular respiration *via* the formation of mitochondria-associated ER membranes (MAMs) (Betz et al., 2013). Hence, we evaluated the effects of sustained glucocorticoid exposure on MAMs by measuring mitochondria-ER colocalization using confocal microscopy. Sustained DEX exposure decreased mitochondria-ER colocalization detected by PC, PC overlap, Mander’s M1, Coste’s PC and Li’s ICA (Figure 7A, 7B), suggesting an overall reduction in MAMs.

**Figure 7.**
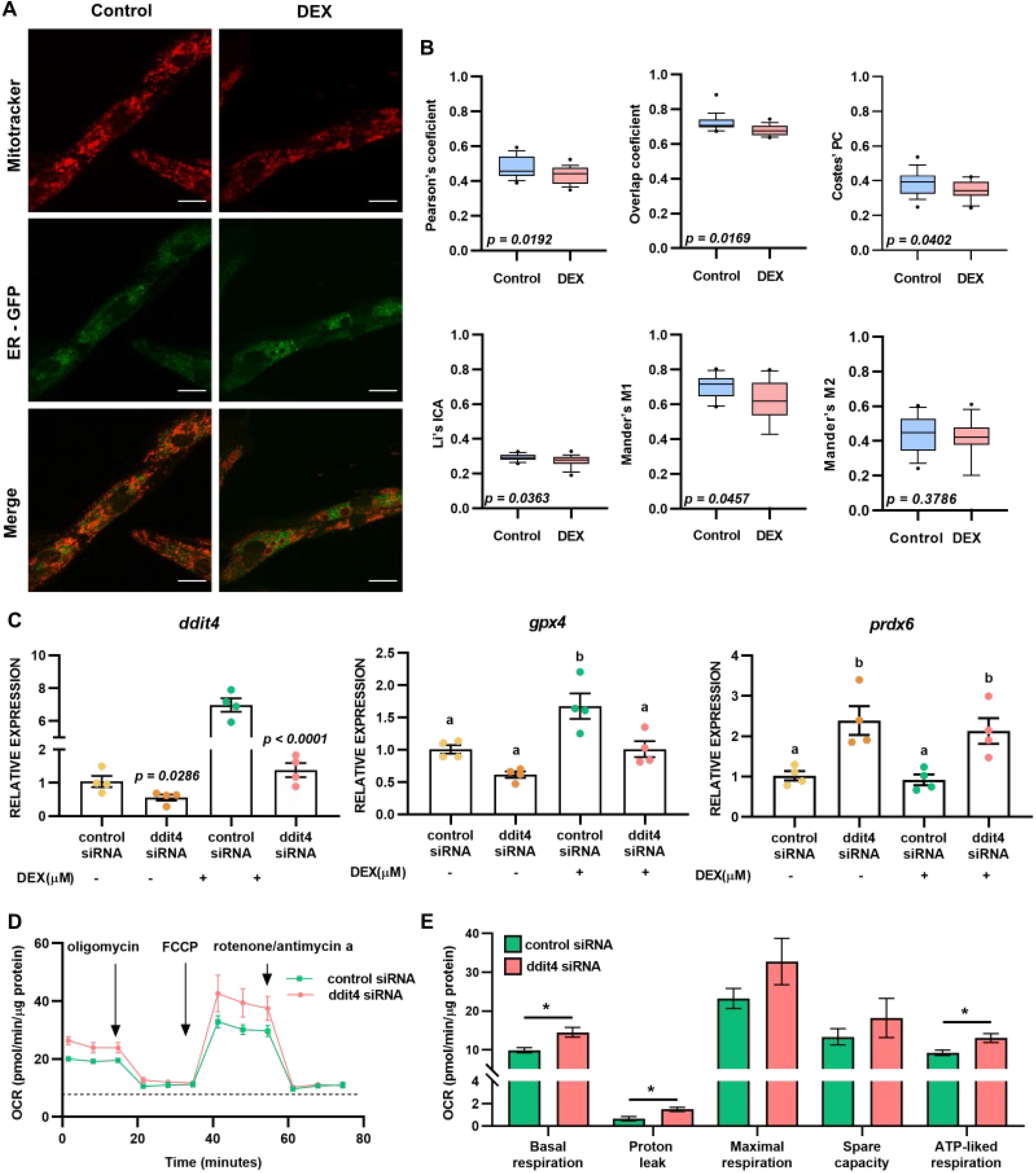
Sustained glucocorticoid exposure disrupts mitochondria-associated ER membranes and decreases mitochondrial function via ddit4. A) Mitochondria (red) and ER (green) colocalization in myotubes treated with or without 100 µM DEX for 48h (scale bar is 21 µm). B) Mitochondria-ER colocalization coefficients. Whiskers show 10-90 percentile. C) Effect of ddit4 silencing on gpx4 (F=12.76, p=0.0005) and prdx6 (F=8.718, p=0.0024) expression. Different letters denote significant differences between treatments. D) Effect of ddit4 knockdown on oxygen consumption and E) mitochondrial function in myotubes treated with DEX. * = p<0.05. Proton leak (p=0.0086), basal (p=0.0116), maximal (p=0.0997) and ATP-linked respiration (p=0.0222). Data are mean ± SEM. n=6.

DNA damage-inducible transcript 4 (DDIT4/REDD1), a shared target of the three TFs identified in our analysis (Figure 3D), is a stress protein that represses mTOR complex 1 during glucocorticoid exposure (Britto et al., 2014; Wang, Kubica, Ellisen, Jefferson, & Kimball, 2006). Recent evidence also suggests that DDIT4 promotes MAMs dissociation (Britto et al., 2018). In our experiments, DEX increased *ddit4* expression *via* GR signaling (Figure 4A, Figure 4D) while reducing MAMs (Figure 7A, 7B) and suppressing mitochondrial metabolism (Figure 5). Therefore, we evaluated the effects of *ddit4* knockdown (Figure 7B) on mitochondrial function in myotubes treated with DEX (Figure 7C). *ddit4* silencing increased basal oxygen consumption, proton leak, and ATP-linked respiration (Figure 7C, 7D, 7E). In contrast, *ddit4* silencing did not have an effect on *gr, foxo1*, or *cebpd* expression (Figure S5), suggesting a direct role for DDIT4 in the suppression of mitochondrial metabolism in myotubes treated with DEX. *ddit4* knockdown also increased the expression of the antioxidant peroxiredoxin 6 (*prdx6*) while decreasing glutathione peroxidase 4 (*gpx4*) expression, suggesting an additional role for DDIT4 in modulating redox homeostasis. Collectively, these results show that sustained glucocorticoid exposure increases *ddit4* expression, disrupts MAMs, promotes mitochondrial fission, and suppresses mitochondrial metabolism. The results also show that DDIT4 directly suppresses mitochondrial respiration and regulates redox metabolism during sustained glucocorticoid exposure. Hence, DDIT4 plays a crucial role in metabolic adaptation to glucocorticoid-induced energetic stress in elephant seal myotubes.

## Discussion

We generated a cellular model to study the effects of sustained glucocorticoid exposure on elephant seal muscle metabolism. Our model retains key physiological components of skeletal muscle, including expression of myogenic genes, the formation of contractile myofibers, and a distinctive metabolic phenotype characterized by increased mitochondrial content and a switch from glycolysis to oxidative phosphorylation as a source of ATP during myogenesis (Kraft et al., 2006; Moyes, Mathieu-Costello, Tsuchiya, Filburn, & Hansford, 1997). Elephant seal myotubes are amenable to pharmacological manipulation, expression of exogenous DNA, and gene silencing. Thus, our model provides a system to study the molecular underpinnings of the extreme adaptations exhibited by elephant seals. Using our model, we show that sustained exposure to glucocorticoids is associated with transcriptional signatures of muscle remodeling and upregulation of genes that promote cell survival in elephant seal myotubes. We also found that adaptation to sustained glucocorticoid exposure involves changes in mitochondrial morphology and function, MAMs dissociation, and a metabolic shift towards glycolysis.

Our RNA-seq analyses identified enriched PI3K-Akt signaling in all DEX treatments. PI3K-Akt signaling inhibits DEX-induced proteolysis in L6 myotubes by inactivating glycogen synthase kinase 3 beta (B. G. Li, Hasselgren, & Fang, 2005). Moreover, PI3K-Akt signaling promotes redox homeostasis by regulating malic enzyme 1 and isocitrate dehydrogenase 1, which are crucial for NADPH production (Hoxhaj & Manning, 2020). Hence, PI3K-Akt signaling likely supports cellular adaptation to sustained glucocorticoid exposure in seal myotubes by inhibiting proteolysis and promoting redox balance. Consistent with this observation, our data show that DEX upregulated the cytoprotective phospholipid hydroperoxidase *gpx4* without increasing the expression of *myostatin* or *murf1*, which mediate DEX-induced muscle wasting in rats, C2C12 and L6 myotubes (Ma et al., 2001; Ma et al., 2003).

PI3K-Akt signaling also promotes cell survival by increasing glucose uptake (DeBerardinis, Lum, Hatzivassiliou, & Thompson, 2008). In our experiments, DEX suppressed mitochondrial respiration without affecting glycolysis-driven ATP production. Glycolysis is less efficient than oxidative phosphorylation but represents a metabolic strategy that promotes cell survival during energetic stress (Epstein, Gatenby, & Brown, 2017; Liberti & Locasale, 2016). Reduced mitochondrial activity is a hallmark of the Warburg effect, which promotes survival by supporting a high rate of glycolysis in both cancer (Seyfried & Shelton, 2010; Vander Heiden, Cantley, & Thompson, 2009), and non-cancer cells (Abdel-Haleem et al., 2017). In our experiments, DEX increased pyruvate dehydrogenase kinase 1 (*PDK1*) expression while decreasing respiratory but not glycolytic capacity. PDK1 promotes energy homeostasis by blocking the incorporation of pyruvate into the TCA cycle in skeletal muscle, favoring the Warburg effect (Zhang, Hulver, McMillan, Cline, & Gilbert, 2014). Thus, our results suggest that adaptation to sustained glucocorticoid exposure in elephant seal myotubes involves a metabolic shift towards glycolysis. In fasting elephant seals, 99% of endogenous glucose production enters glycolysis but only 16% of this carbon is committed to the TCA cycle and oxidation, with the bulk of the pyruvate being recycled to glucose through lactate gluconeogenesis (Champagne et al., 2012; Houser, Crocker, Tift, & Champagne, 2012; Tavoni, Champagne, Houser, & Crocker, 2013).

Decreased mitochondrial function in myotubes treated with DEX was associated with increased mitochondrial fragmentation. Dysregulated mitochondrial dynamics (fission/fusion events) alters mitochondrial function in skeletal muscle (Sebastián et al., 2016; Tezze et al., 2017; Whitley, Engelhart, & Hoppins, 2019). In our studies, DEX increased DRP1 phosphorylation while promoting mitochondrial fission. DRP1 drives the effect of glucocorticoids on mitochondrial fragmentation in hepatoma cells (Hernández-Alvarez et al., 2013), but DRP1 knockdown or muscle-specific DRP1 deletion induces muscle atrophy and mitochondrial dysfunction in mice (Dulac et al., 2020; Favaro et al., 2019). Mitochondrial fission occurs upon DRP1 recruitment at positions where ER tubules contact mitochondria (Friedman et al., 2011). These mitochondria-ER contact sites are spatially and functionally connected by MAMs, which participate in calcium transport between both organelles, inflammation, bioenergetics, cell death, lipid biosynthesis, and autophagy (Missiroli et al., 2018; Sassano, van Vliet, & Agostinis, 2017). Chronic MAMs disruption impairs cardiomyocyte and hepatic glucose sensing and dysregulates mitochondrial dynamics and function (Gutiérrez et al., 2014; Theurey & Rieusset, 2017; Theurey et al., 2016). Our data show that DEX induces MAMs disruption and reduces mitochondrial function in seal myotubes. Intriguingly, recent evidence suggests that MAMs disruption is an adaptive mechanism that might promote cell survival during glucocorticoid-induced energetic stress by suppressing mitochondrial respiration, resulting in a low energy consumption state (Britto et al., 2018).

The proposed mechanism for MAMs disruption involves DDIT4/REDD1 binding to glucose-regulated protein 75, which is responsible for mitochondria-ER coupling (Britto et al., 2018; Honrath et al., 2017; Szabadkai et al., 2006). DDIT4 deletion prevents DEX-induced skeletal muscle atrophy (Britto et al., 2014) and increases oxygen consumption in permeabilized mouse skeletal muscle fibers, whereas DDIT4 knockdown prevents interaction of MAMs proteins in human myoblasts (Britto et al., 2018). Our studies show that DEX promotes MAMs dissociation and suppresses mitochondrial respiration while increasing GR-dependent *ddit4* expression. Moreover, our results also show that *ddit4* knockdown recovers mitochondrial respiration in myotubes treated with DEX. Although we did not directly evaluate the effect of DDIT4 on mitochondrial morphology or MAMs, our functional data suggest that DDIT4 promotes MAMs dissociation in seal myotubes exposed to DEX. Moreover, our results also show that DDIT4 is a shared target of CEBPD, CEBPB and MTF1, the main transcriptional regulators of the sustained response to glucocorticoids in seal myotubes. DDIT4 is also one of the top skeletal muscle genes upregulated after ACTH administration in elephant seals (Khudyakov, Champagne, et al., 2015).

Our results also show that besides suppressing mitochondrial metabolism, DDIT4 regulates *prdx6* expression. Prdx6 supports muscle function (Pacifici et al., 2020; Sakellariou et al., 2018) by limiting lipid peroxidation (Arevalo & Vázquez-Medina, 2018; Fisher et al., 2018). Growing evidence shows that DDIT4 regulates redox homeostasis, but its specific role in oxidant generation or removal remains unclear. While loss of *ddit4* in mouse embryonic fibroblasts increases mitochondrial superoxide and hydrogen peroxide production (Horak et al., 2010), *ddit4* knockdown upregulates superoxide dismutase, catalase, and glutathione peroxidase in human endothelial cells exposed to endotoxin (Hou, Yang, & Yin, 2019). Moreover, induction of a DDIT4/thioredoxin-interacting protein complex, which increases cellular oxidant levels by suppressing thioredoxin expression and activity (Patwari, Higgins, Chutkow, Yoshioka, & Lee, 2006), is necessary for stress-induced autophagy (Alvarez-Garcia et al., 2017; Qiao et al., 2015). Thus, our results show that DDIT4 plays a crucial role in cellular adaptation to sustained glucocorticoid exposure in elephant seal myotubes by regulating mitochondrial metabolism and redox homeostasis. Furthermore, our results suggest that DDIT4 may play a role in the extraordinary oxidative stress tolerance exhibited by elephant seals undergoing prolonged food deprivation, which increases circulating glucocorticoid levels (Vázquez-Medina, Crocker, Forman, & Ortiz, 2010; Vázquez-Medina et al., 2013; Vázquez-Medina, Zenteno-Savín, Forman, Crocker, & Ortiz, 2011).

In summary, here we show that adaptation to sustained glucocorticoid exposure in elephant seal myotubes involves concerted changes in the expression of genes that promote cell survival and muscle remodeling. We also show that sustained exposure to glucocorticoids causes a metabolic shift towards glycolysis, which is supported by reduced mitochondrial function and changes in mitochondrial morphology and decreased mitochondria-ER interactions. Furthermore, our results show that DDIT4 plays a critical role in adaptation to sustained glucocorticoid exposure in elephant seal myotubes by modulating mitochondrial function and redox homeostasis.

## Acknowledgments

JMT-V was supported by a UC MEXUS-CONACyT postdoctoral fellowship. Confocal imaging was conducted at the UC Berkeley CRL Molecular Imaging Center, supported by NSF DBI-1041078. We thank Holly Aaron and Feather Ives for their microscopy training and assistance, and Kaitlin Allen and Emily Lam for their support in the field. Library preparation and sequencing were conducted at UC Berkeley’s Functional Genomics and Vincent J. Coates Genomics Sequencing laboratories. Research was funded by UC Berkeley startup funds.

## Methods

### Animals

Animal handling protocols were approved by Sonoma State University and UC Berkeley Institutional Animal Care and Use Committees. Sampling was conducted under National Marine Fisheries Service Permit (NMFS) No. 19108. Cell lines were developed under NMFS Permit No. 22479. Juvenile (∼10 month old) northern elephant seals (*Mirounga angustirostris*) were sampled at Año Nuevo State Reserve (San Mateo County, CA). Animals were chemically immobilized using tiletamine-zolazepam and ketamine as previously described (Vázquez-Medina et al., 2010). Muscle biopsies were collected from the posterior flank region of each animal using a 6.0 mm diameter biopsy punch (Vázquez-Medina et al., 2010). Samples were rinsed with ice-cold HBSS (Gibco, Thermo Fisher Scientific, Waltham, MA), placed in Ham’s F-10 nutrient mix (Gibco) containing penicillin-streptomycin (Gibco), and transported on ice to the laboratory.

### Isolation, maintenance and differentiation of primary cells

Tissues were minced into small pieces and incubated with collagenase (Wortington, Danvers, MA). Cell suspensions were passed through 18g and 22g syringes to dissociate tissue. Collagenase was neutralized with two volumes of complete growth medium: Ham’s F-10 (Gibco) supplemented with fetal bovine serum (Seradigm, VWR, Radnor, PA), HEPES (Gibco) and Antimycotic/Antibiotic solution (Gibco). Cell suspensions were filtered using a 100 µm strainer and centrifuged. Cell pellets were resuspended in growth medium and plated in tissue culture dishes coated with type I rat tail collagen (Sigma, St. Louis, MO). Cultures were expanded and frozen at passage 1. Myoblasts were enriched by pre-plating, and differentiated into myotubes by incubation with DMEM (Gibco) supplemented with 5% horse serum (Gibco) for seven days. The phenotype of the preparation was confirmed by immunostaining with desmin antibodies and RT-qPCR for myogenic genes.

### Dexamethasone treatment

Myotubes were treated with four ascending concentrations (0.1, 1, 10 and 100 µM) of the glucocorticoid dexamethasone (DEX) (BioVision, Milpitas, CA) for 48 hrs. Cells were harvested for RNA or protein extraction, or fixed for confocal imaging after treatment. In some experiments, cells were co-treated with the glucocorticoid receptor antagonist RU486 (10 and 100 µM). Live cells were used for bioenergetic studies.

### RNA extraction and RT-qPCR

Total RNA was extracted using Trizol (Invitrogen, Carlsbad, CA). Genomic DNA contamination was removed using a Turbo DNA-free kit (Thermo Fisher), and confirmed by lack of genomic DNA amplification using RT-PCR. RNA was quantified with a Qubit fluorometer (Invitrogen). cDNA was synthesized using a High-Capacity cDNA Reverse Transcription kit (Invitrogen). qPCR was performed with DyNAmo Flash SYBR Green Master Mix (Thermo Fisher) under the following conditions: 7 min at 95°C, followed by 40 cycles of 20 s at 95 °C, and 30 s at 60°C. Relative expression was calculated using the comparative 2^-ΔΔCT^ method with the geometric mean of *gapdh* and *actin* (Livak & Schmittgen, 2001). Primer sequences are listed in Supplementary Table 1.

### RNA-seq

RNA integrity was measured using an Agilent 2100 Bioanalyzer (Agilent Technologies, Santa Clara, CA). RNA with RIN > 9.5 was used for library preparation and sequencing at UC Berkeley’s Functional Genomics and Vincent J. Coates Genomics Sequencing Laboratories. cDNA libraries were prepared from poly(A)-captured mRNA using a KAPA RNA HyperPrep Kit (Roche, Basel, Switzerland, Cat No. KK8541) with Truseq adapters. Libraries were generated from three replicates for each treatment, and sequenced to a total of 25 million reads per sample on a Novaseq platform (Illumina, San Diego, CA).

### Genome annotation

Annotation of the HiC scaffolded elephant seal genome (https://www.dnazoo.org/assemblies/Mirounga_angustirostris) was conducted using *ab initio* methods and RNA-seq data. Transcripts were *de novo* assembled using Trinity V2.8.4. Open reading frames were extracted using six-frame translation. Potential coding sequences were translated using TransDecoder V5.5.0 (Haas, BJ, https://github.com/TransDecoder/TransDecoder). The longest transcript isoforms (336,657) were mapped to the genome using Minimap2 (H. Li, 2018) with splice-aware parameters (splice:hq). Protein coding sequences (181,409) were mapped with SPALN V2.3.3 (Iwata & Gotoh, 2012) using tetrapod-specific parameters. Additional evidence on gene boundaries was generated from RNA-seq alignments to the genomic reference using STAR aligner (Dobin et al., 2013). Genomic repeats were softmasked with RepeatMasker V4.0.9 (Tarailo-Graovac & Chen, 2009) using the mammalian-specific database. This repeat-masked genome was annotated *ab initio* with BRAKER2 (Hoff, Lomsadze, Borodovsky, & Stanke, 2019). The alignments from RNA-seq data, transcripts, and proteins were used along with the *ab initio* models to generate evidence-weighted gene predictions using Funannotate V1.7.4 (Jon Love et al., 2020, Funannotate v1.7.4. Zenodo. https://doi.org/10.5281/zenodo.3679386). 58,999 genes were predicted and filtered based on similarity search against a database of vertebrate proteins from ENSEMBL 99 (Yates et al., 2020), retaining a total of 26,745 gene models.

### Transcriptome analyses

Transcript levels were quantified using RSEM 1.3.1 (B. Li & Dewey, 2011). Genes differentially expressed (DE) between control and each DEX treatment, as well as between successive treatments, were identified at an FDR of 5% using EBSeq. Expression levels were converted to z-scores normalized by the mean and variance of each gene across all conditions and replicates. For each condition, the median z-score was selected for each gene and a Euclidean distance matrix was constructed. Genes that changed in a concerted manner with increasing DEX concentrations were clustered using k-means. Diminishing returns were observed after 6 clusters, thus k = 6 was selected as the optimal clustering. Gene set enrichment analysis with gene ontology (GO) terms was used to identify biological processes enriched in the gene clusters. Functional interaction networks (FIN) were built from enriched Reactome pathways using Cytoscape v3.8.1 (Wu et al., 2010; Wu & Haw, 2017). Cis-regulatory analysis was performed using iRegulon in Cytoscape (Janky et al., 2014).

### Gene silencing

Elephant seal-specific siRNAs were designed using IDT’s RNAi design tool (IDT Corporation, Newark, NJ). Myotubes were transfected with siRNAs targeting glucocorticoid receptor (*gr*), DNA damage inducible transcript 4 (*ddit4*), or with non-targeting sequences (control siRNA) using Lipofectamine RNAiMAX (Thermo Fisher). Gene knockdown was confirmed 72h post-transfection using RT-qPCR. siRNA sequences are shown in Supplementary Table 2.

### Confocal imaging

Myotubes grown in glass-bottom chambers were transfected with CellLight ER-GFP BacMam 2.0 fusion constructs (Thermo Fisher) following the manufacturer’s instructions. Mitochondria were labeled with MitoTracker Red-CMXRos (Cell Signaling Technology, Danvers, MA). Cells were fixed with ice-cold methanol, rinsed with PBS, and imaged in VECTASHIELD mounting media (Vector Laboratories, Burlingame, CA) using a Zeiss LSM 710 microscope fitted with 40X (Plan-Apochromat 1.4 NA oil, 0.13mm) and 63X (Plan-Apochromat 1.4 NA oil, 0.19 mm) objectives. Visualization of mitochondrial networks and mitochondria-ER colocalization was conducted at 40X and 63X, respectively. Magnification was set to 1 x zoom (pixel size = 0.4151) for 40X and 1 x zoom (pixel size = 0.2636) for 63X. Z-slices were collected at 1 µm for 40X and 0.5 µm for 63X.

### Analyses of mitochondria-ER interactions

Mitochondria-ER colocalization analyses were conducted using Fiji with the JACoP v2.0 plugin (Bolte & Cordelières, 2006; Cordelieres & Bolte, 2008). Mitochondria-ER interactions were evaluated using Pearson’s correlation coefficient (PC), PC overlap coefficient, Mander’s colocalization coefficients, Costes’ PC (Costes et al., 2004), and Li’s intensity correlation analysis (ICA) (Q. Li et al., 2004). PC and PC overlap estimate the association between two objects in a dual-channel. Mander’s M1 and M2 represent colocalization of one channel over the other, where M1 is the proportion of pixels in mitochondria concurrent with pixels in the ER over its total intensity, and M2 is the proportion of pixels in the ER concurrent with pixels in mitochondria over its total intensity (Manders, Verbeek, & Aten, 1993). Costes’ PC indicates the statistical estimation of colocalization due to co-compartmentalization or interaction, excluding random overlap after 200 randomized tests (Costes et al., 2004). Li’s ICA normalizes the intensity per channel, and correlates the intensity distribution of each channel with the covariance of both channels (Bolte & Cordelières, 2006; Cordelieres & Bolte, 2008; Q. Li et al., 2004).

### Mitochondrial network analyses

Mitochondrial network analyses were conducted according to (Harwig et al., 2018). Raw images were processed with mean filtering, contrast limited adaptive histogram equalization, and enhanced contrast in Fiji (Valente, Maddalena, Robb, Moradi, & Stuart, 2017), and analyzed using MitoGraph v3.0 (https://github.com/vianamp/MitoGraph). MitoGraph creates skeletonized representations of the mitochondrial network, which is subdivided into edges (mitochondrial tubules). Edges are joined at nodes that represent the end of mitochondria and branch points between mitochondrial interconnections. MitoGraph also measures mitochondrial volume (voxels), length, and relative width. MitoGraph-generated data were processed using R-scripts (https://github.com/Hill-Lab/MitoGraph-Contrib-RScripts). We obtained total edges, nodes, length, number of connected components, relative size of the largest connected component to total mitochondrial size (PHI), average edge length (Avg), a connectivity score indicative of the sum of values increased in the fusion state (PHI, Avg length and Avg degree) divided by the sum of values increased in the fission state (total nodes, edges and connected components), and a degree of distribution (1, 2, 3, 4, and AVG), which indicates free ends or number of junctions. Values were normalized to total mitochondrial length to control for myotube size. ParaView was used to generate 3D representations.

### Immunofluorescence

Immunofluorescence was conducted as previously described (Vázquez-Medina et al., 2016). Cells were fixed with ice-cold 1:1 acetone/methanol, permeabilized, blocked, and incubated overnight with desmin (Thermo Fisher, PA5-17182) or myosin heavy chain 1 (R&D systems, Minneapolis, MN, MAB4470) antibodies diluted 1:100. Alexa Fluor secondary antibodies (Thermo Fisher) were used at a 1:200 dilution. Samples mounted in VECTASHIELD media (Vector Laboratories) were imaged using Zeiss LSM 710 (MYH1) or LSM 780 (desmin) confocal microscopes fitted with 40X objectives and Zen software.

### Western blot

Immunoblotting was conducted as previously described (Vázquez-Medina et al., 2019). Cells were sonicated in a phosphate buffer containing protease and phosphatase inhibitors. Proteins were resolved by SDS-PAGE, transferred onto nitrocellulose membranes, incubated with Revert 700CW total protein stain (Licor, Omaha, NE) and imaged using a two-color near-infrared system (Azure c500, Azure Biosystems, Dublin, CA). Membranes were blocked with Odyssey blocking buffer (Licor), and incubated overnight with the following primary antibodies diluted 1:1,000: COX4 (Novus Biologicals, Littleton, CO, NB110-39115), GR (Invitrogen, PA1-511A), DRP1 (Cell Signaling Technology, 8570), phospho-DRP1Ser616 (Cell Signaling Technology, 3455), and HPRT (Novus Biologicals, NBP2-75528). IRDye 800CW secondary antibodies (Licor) were used at a 1:4,000 dilution. Individual bands were quantified using Fiji. Density values were normalized to total protein or HPRT.

### Extracellular flux assays

Cells seeded in Seahorse plates (Agilent Technologies) were washed with assay medium (Seahorse XF DMEM supplemented with 6mM glucose, pH 7.4) and placed in a CO_2_-free incubator for 1 h. Oxygen consumption (OCR) and extracellular acidification rates (ECAR) were measured in a calibrated Seahorse XFp extracellular flux analyzer (Agilent) in the presence of oligomycin (1 µM), carbonyl cyanide-p-trifluoromethoxyphenylhydrazone (FCCP, 2 µM), and rotenone/antimycin A (0.5 µM). Values were normalized to cell number. Mitochondrial function was calculated according to (Divakaruni, Paradyse, Ferrick, Murphy, & Jastroch, 2014). Cellular ATP production was calculated according to (Mookerjee et al., 2018). Protein concentration was measured using a Qubit protein assay (Invitrogen).

### Statistical analyses

Normality and homoscedasticity were evaluated using D’Agostino-Pearson and Bartlett’s tests. Differences among groups were determined using t-tests, 1- or 2-way ANOVAs followed by Fisher’s post hoc analyses. Non-parametric data were evaluated using Mann-Whitney or Kruskal-Wallis tests with Dunn’s post hoc analyses. Differences were considered significant when *p* < 0.05. Statistical analyses were conducted using GraphPad Prism v9. Results are presented as mean ± SEM.

