## Supplementary Tables and Figures for "Elephant seal muscle cells adapt to sustained glucocorticoid exposure by shifting their metabolic phenotype"

### Supplementary material

#### *Supplementary Table 1. Primer sequences*

| *Gene* | *Sequence 5 ' to 3'* |
|---|---|
| ***gapdh*** | *CAA GGC TGA GAA CGG GAA GC* |
| | *ATC GGC AGA GGG AGC AGA GA* |
| ***actin*** | *CGG TCA GTT CAT GGC TGA GG* |
| | *AAG GCT CGG ACC TTC CCA AC* |
| ***cebpd*** | *AAC CCA CAC CAC CCA CGT C* |
| | *TCC ACC AGC TTC TGC TGC AT* |
| ***ddit4*** | *ACG TCT GCG TGG AGC AAG G* |
| | *GTG CTC AGC GTC AGG GAC TG* |
| ***foxo1*** | *GCA CCG GGA CTC TTG AAG GA* |
| | *GTG TGG GAG CTG GGG TTC AT* |
| ***foxo3*** | *TCT GCG GGG TGG AAG AAC TC* |
| | *GCA GGG CTG CCT TCT TCT TG* |
| ***mstn*** | *TGG GCT TGA TTG TGA TGA GCA* |
| | *AGG GGC CTG CTG AAC CTC TT* |
| ***mrf4*** | *GCT GCC CAA GGT GGA GAT TC* |
| | *TTA TCA CGA GCC CCC TGG AA* |
| ***mtor*** | *ACC AGC CGA TCA TTC GCA TT* |
| | *GGG TGT TCA CCA GGC CAA AG* |
| ***murf1*** | *CCG TCA CGA GGT GAT CAT GG* |
| | *AGC ACG TGG GCA TCT CAC AC* |
| ***myod*** | *ACT GCC TGT CCA GCA TCG TG* |
| | *TAG ATC GGG TTG GGG TTT GC* |
| ***myog*** | *AGG CTA CGA GCG GAC TGA GC* |
| | *CCT TCT TGA GCC TGC GCT TC* |
| ***pcna*** | *TGG ACT CCT CCC ACG TCT CC* |
| | *GCC AGG GTG TCC GCA TTA TC* |
| ***gr*** | *AGC AGG CCA CTC CAG GAACG* |
| | *AGT GTG GTC ATG ATC CGC CAA GT* |

| | |
|---|---|
| ***myh1*** | *TTG ACG ACC TTG AGC TGA CC* |
| | *CTG CCA TCT CTT CCG TGA GG* |
| ***myh8*** | *GAG ACA AGC TGA GGA GGC TG* |
| | *GTT GAC CTG GGA CTC AGC AA* |
| ***tnnt1*** | *TCC GCT GTT CAA AAT GCA CG* |
| | *ATT CCC CCG AAG ATC CCA GA* |
| ***mr*** | CCA GCA ATG TGG GCT CTC CT |
| | TTT GTG TTG GAA GGG CTG GAA |

### *Supplementary Table 2. siRNA sequences*

| Gene | Sequence 5 ' to 3' | |
|---|---|---|
| ***gr*** | Sense | rGrArGrArUrCrUrGrArCrCrUrCrUrUrGrArUrArGrArUrGAA |
| | Antisense | rUrUrCrArUrCrUrArUrCrArArGrArGrGrUrCrArGrArUrCrUrCrCrA |
| ***ddit4*** | Sense | rGrGrGrCrUrUrCrCrGrArGrUrCrArUrCrArArGrArArGrAAG |
| | Antisense | rCrUrUrCrUrUrCrUrUrGrArUrGrArCrUrCrGrGrArArGrCrCrCrGrU |

**Supplementary Table 3. DE genes involved in the Central Carbon Metabolism in Cancer pathway.**

| *Gene* | | *PPEE* | *100 µM DEX* vs *control* |
|---|---|---|---|
| *BHLHE40*† | MYC proto-oncogene, bHLH transcription factor | $6.91E^{-06}$ | DOWN |
| *EGFR*† | Epidermal growth factor receptor | $7.84E^{-03}$ | UP |
| *ERBB3* * | Receptor tyrosine-protein kinase erbB-3 | $7.83E^{-03}$ | DOWN |
| *HIF1A*† | Hypoxia inducible factor 1 subunit alpha | $0.00E^{+00}$ | DOWN |
| *HK2*† | Hexokinase 2 | $1.54E^{-02}$ | DOWN |
| *IDH1* * | Isocitrate dehydrogenase 1 (NADP+), soluble | $3.33E^{-16}$ | UP |
| *MAP2K1* * | Mitogen-activated Protein Kinase Kinase 1 | $2.11E^{-02}$ | DOWN |
| *MAP3K10* * | Mitogen-Activated Protein Kinase Kinase Kinase 10 | $8.15E^{-05}$ | UP |
| *MAP3K12* * | Mitogen-Activated Protein Kinase Kinase Kinase 12 | $1.09E^{-06}$ | DOWN |
| *MAP3K3* * | Mitogen-Activated Protein Kinase Kinase Kinase 3 | $7.33E^{-05}$ | UP |
| *MAP3K4* * | Mitogen-Activated Protein Kinase Kinase Kinase 4 | $4.64E^{-02}$ | DOWN |
| *MAP3K7* * | Mitogen-Activated Protein Kinase Kinase Kinase 7 | $1.13E^{-06}$ | DOWN |
| *MAP4K1* * | Mitogen-Activated Protein Kinase Kinase Kinase Kinase 1 | $3.81E^{-05}$ | DOWN |
| *MAP4K5* * | Mitogen-Activated Protein Kinase Kinase Kinase Kinase 5 | $1.99E^{-02}$ | DOWN |
| *MAPK8IP3** | Mitogen-activated protein kinase 8 interacting protein 3 | $5.24E^{-01}$ | UP |
| *MTOR* * | Mechanistic target of rapamycin kinase | $2.32E^{-02}$ | UP |
| *PDK1*† | Pyruvate dehydrogenase kinase 1 | $2.21E^{-06}$ | UP |
| *PDK2*† | Pyruvate dehydrogenase lipoamide kinase isozyme 2 | $1.34E^{-06}$ | UP |
| *PDK4*† | Pyruvate dehydrogenase lipoamide kinase isozyme 4 | $0.00E^{+00}$ | UP |
| *PFKFB4*† | 6-phosphofructo-2-kinase/fructose-2,6-bisphosphatase 4 | $1.10E^{-09}$ | UP |
| *PFKL*† | 6-phosphofructokinase, liver type | $1.24E^{-12}$ | DOWN |
| *PGAM1*† | Phosphoglycerate Mutase 1 | $0.00E^{+00}$ | DOWN |
| *PIK3R1*† | Phosphoinositide-3-kinase regulatory subunit 1 | $0.00E^{+00}$ | UP |
| *PIK3R2*† | Phosphoinositide-3-kinase regulatory subunit 2 | $1.47E^{-02}$ | DOWN |
| *PIK3R5*† | Phosphoinositide-3-kinase regulatory subunit 5 | $2.22E^{-15}$ | UP |
| *RASL11B*† | RAS-like, family 11, member B | $3.28E^{-03}$ | UP |
| *RASSF4* * | Ras association domain family member 4 | $3.21E^{-06}$ | DOWN |

| | | | |
|---|---|---|---|
| *SIRT7*† | Sirtuin 7 | 2.66E-15 | UP |
| *SLC1A5*† | Solute carrier family 1 (neutral amino acid transporter), member 5 | 5.37E-08 | UP |
| *SLC7A5*† | Solute carrier family 7 (neutral amino acid transporter), member 5 | 1.36E-03 | DOWN |
| *TP53*† | Tumor protein p53 | 1.29E-07 | DOWN |
| *TP53BP1** | TP53-binding protein 1 | 1.27E-02 | DOWN |

*Genes DE only in control vs DEX 100 μM †Genes DE in other DEX treatments

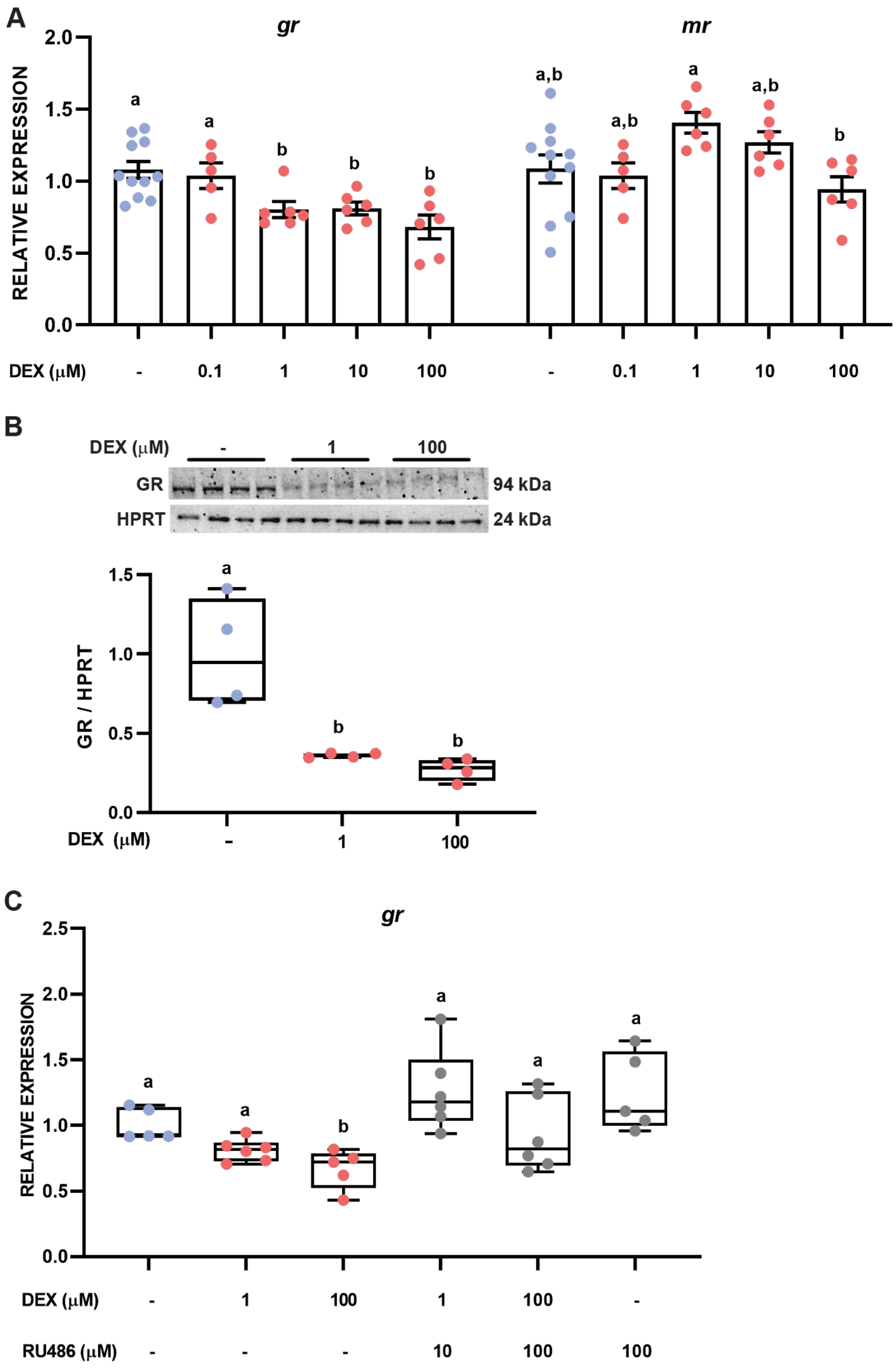


***Supplementary Figure 1. Dexamethasone signals via glucocorticoid receptors in elephant seal myotubes.*** *A) Changes in mRNA levels of gr (F=4.651, p=0.0055) and mr (F=3.506, p=0.0188) in myotubes treated with DEX for 48h. B) GR protein abundance in myotubes treated with DEX for 48h (p=0.0013, F=15.37). Values are normalized to HPRT. C) gr mRNA levels in myotubes treated with DEX for 48h with or without pre-treatment with the glucocorticoid receptor antagonist RU486 (F=6.805, p=0.0003). Different letters denote significant differences.*

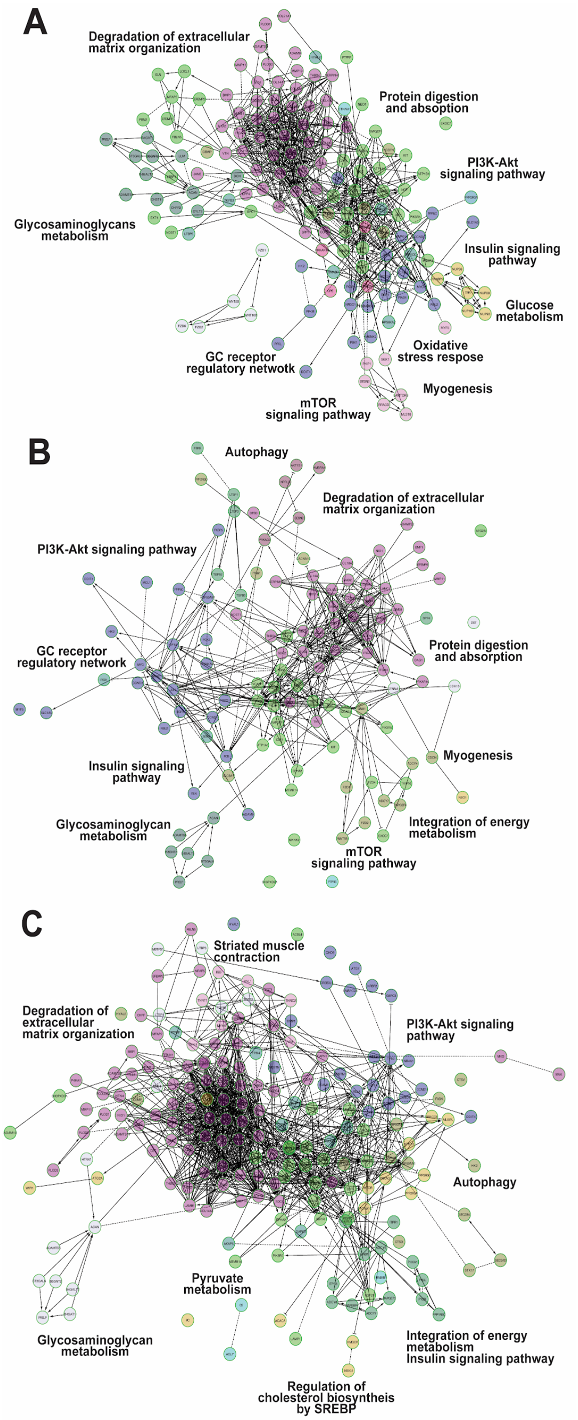


***Supplementary Figure 2. Functional interaction networks for enriched Reactome pathways in myotubes treated with DEX.*** *FIN for DE genes in control vs A) 0.1 µM DEX, B) 1 µM DEX and C) 100 µM DEX.*

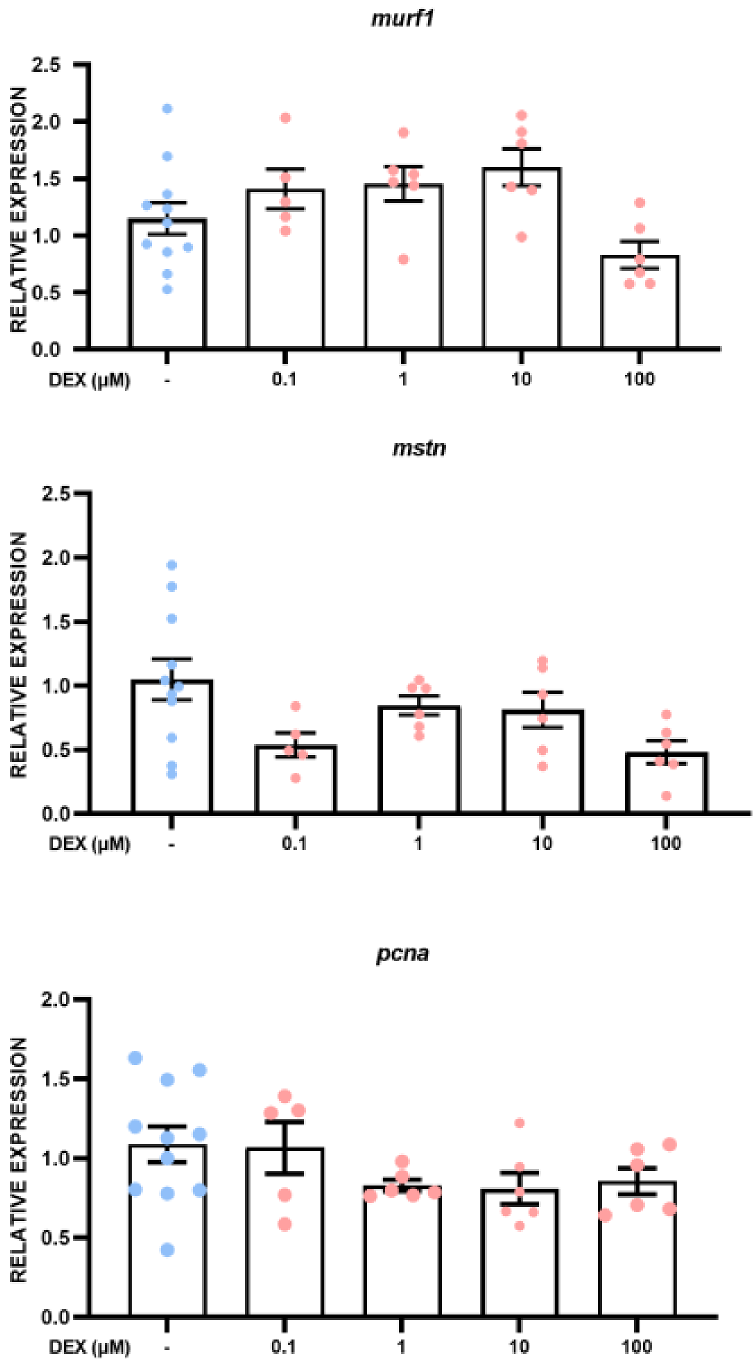


***Supplementary Figure 3. Effect of dexamethasone on the expression of genes related to muscle wasting.*** *Results are mean ± SEM. murf1 (F=3.567, p=0.0175, control vs 10 µM p(adj.)=0.0354), mstn (F=2.999, p=0.0347, control vs 0.1 µM p(adj.)= 0.0161, control vs 100 µM p(adj.)= 0.0053), and pcna (F=1.577, p=0.2068).*

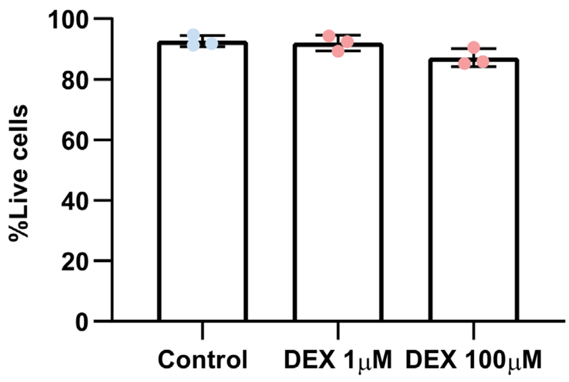


***Supplementary Figure 4. Effects of sustained glucocorticoid exposure on cell viability in elephant seal myotubes.*** *Cell viability was measured in myotubes treated with or without 1 μM or 100 μM DEX. Viability/cytotoxicity was measured using a commercial assay (Thermo Fisher, L3224).* KW=4.356, *p=0.1321*.

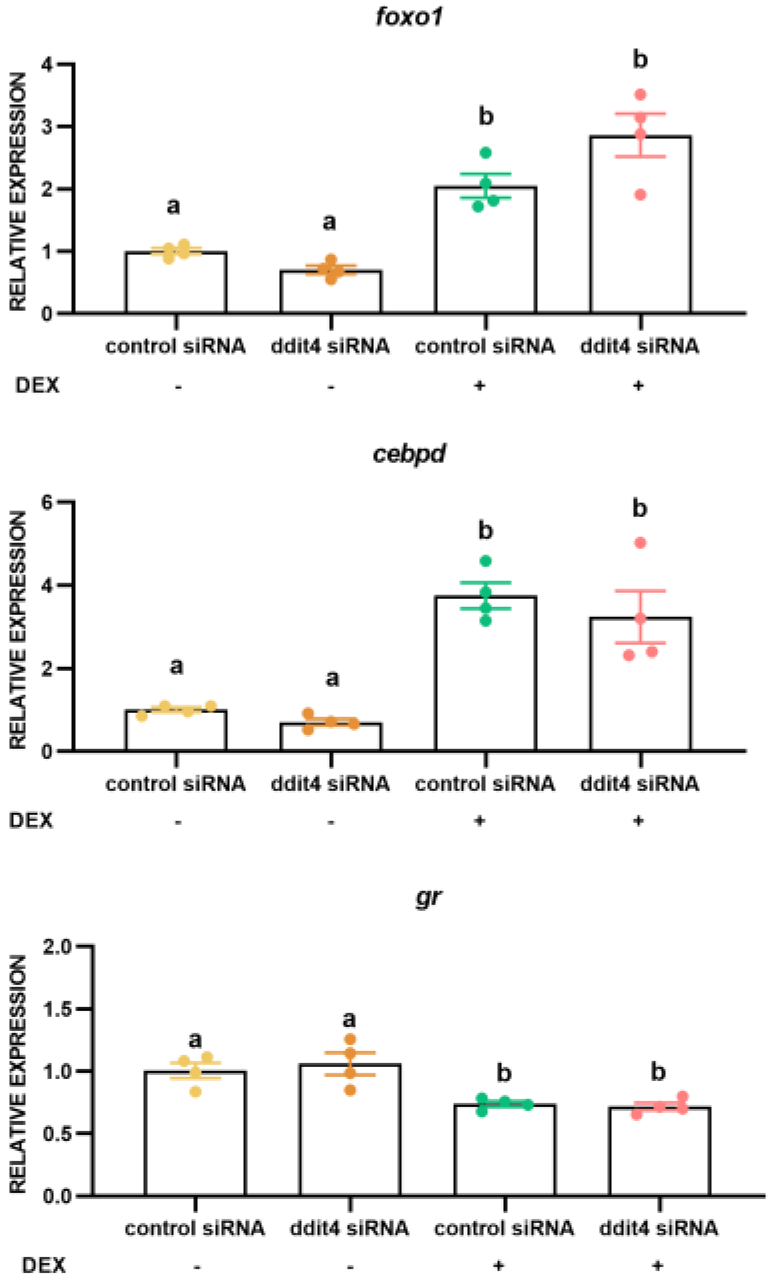


***Supplementary Figure 5. Effect of ddit4 knockdown on* foxo1, cebpd *and* gr *expression in myotubes treated with or without 100 µM DEX for 48h***. *Results are mean* ± *SEM. Different letters denote significant differences between treatments. foxo1 (F=24.29, p<0.0001), cebpd (F=19.02, p<0.0001)*, and *gr (F=9.545, p=0.0017)*.
